# Centriolar satellites regulate *CEP350* mRNA localization and centrosome amplification

**DOI:** 10.64898/2026.03.26.714479

**Authors:** Abraham Martinez, Chad G. Pearson

## Abstract

Messenger RNAs (mRNAs) accumulate at centrosomes in mitosis and interphase, yet the mechanisms governing their localization and the functional significance of centrosomal localization remain poorly understood. Here, we investigate the regulation and function of the centrosome-localized mRNA, *CEP350*. We find that *CEP350* mRNA localizes to centrosomes during S phase via the centriolar satellite protein CEP131 and the RNA binding protein (RBP) Unkempt (UNK), in a microtubule (MT)-dependent manner. CEP131 and UNK stabilize *CEP350* mRNA to maintain *CEP350* mRNA steady-state levels. Furthermore, CEP131 and UNK promote normal CEP350 protein levels at centrosomes. CEP350 is required for PLK4-induced centriole overduplication but is less important for canonical centriole duplication. Moreover, CEP131, UNK, and CEP350 are important for centrosome amplification in triple-negative breast cancer cells. Together, these findings reveal a centriolar satellite-RBP pathway regulating *CEP350* mRNA localization to centrosomes.

**Significance Statement:**

- mRNAs encoding centrosomal proteins localize to centrosomes during interphase, though the mechanism and functional significance of this localization remains unclear.
- This study finds that the centriolar satellite proteins CEP131 and UNK regulate centrosomal localization of *CEP350* mRNA and protein during S phase.
- CEP131, UNK, and CEP350 are required for centrosome amplification in triple negative breast cancer cells – a dependency that identifies them as potential therapeutic targets.

## INTRODUCTION

Translation of localized messenger RNA (mRNA) is a fundamental cellular process enabling protein production precisely where it is needed (Huber et al., 2000; Kang and Schuman, 1996; Martin et al., 1997). Spatial organization of translation is best understood in the context of neurons where synapses are strengthened and weakened, in part, by local translation in a process called synaptic plasticity (Batista et al., 2017; Grzejda et al., 2025; Huber et al., 2000; Kang and Schuman, 1996; Martin et al., 1997). In this context, mRNAs are trafficked to synapses, where they are poised for local translation in response to stimuli, thereby bypassing long-range axonal transport of proteins (Eberwine et al., 2001). Although localized translation has been extensively studied in neurons, somatic cycling cells also locally translate mRNAs at organelles (Kuzniewska et al., 2020; Lécuyer et al., 2007; Pyhtila et al., 2008; Raff et al., 1990; Rangaraju et al., 2019; Safieddine et al., 2021; Sepulveda et al., 2018; Voigt et al., 2017).

Organelles in cycling cells, including mitochondria and the endoplasmic reticulum (ER), reorganize through the cell cycle (Chatre and Ricchetti, 2013; Wang et al., 2013). This reorganization is important at the G1-S phase transition when mitochondrial biogenesis and ER remodeling are required to meet the energetic and protein synthesis demands of cell division (Merta et al., 2021; Mitra et al., 2009). Further, the ER expands throughout S phase, increasing fatty acid biosynthesis (Chowdhury et al., 2024). The reorganization of these organelles requires the localization and local translation of mRNAs (Rangaraju et al., 2019; Reid and Nicchitta, 2015). Recent work suggests mRNAs are translated locally at centrosomes during mitosis to facilitate mitotic spindle organization and chromosome segregation (Groisman et al., 2000; Huang et al., 2015; Ryder et al., 2020; Safieddine et al., 2021; Sharp et al., 2020). How mRNAs are localized and locally translated to impact organelle assembly, reorganization, and function remains poorly understood.

Like mitochondria and the ER, mRNAs localize to centrosomes during interphase, co-localizing with their encoded proteins (Ryder et al., 2020; Sepulveda et al., 2018; Zein-Sabatto et al., 2024). The significance of this interphase mRNA localization remains unclear. The proteins whose mRNAs localize to centrosomes during interphase regulate centrosome-related processes, including centrosome assembly (Safieddine et al., 2021). Centrosome assembly is coordinated with DNA synthesis (Hinchcliffe et al., 1998; Nigg and Holland, 2018). In G1, the centrosome is composed of two microtubule (MT)-based centrioles. At the G1-S phase transition, Polo-Like Kinase 4 (PLK4) is recruited to the proximal ends of both centrioles (Bettencourt-Dias et al., 2005; Habedanck et al., 2005). PLK4 phosphorylates centriole scaffold proteins to initiate the assembly of a new daughter centriole starting at the centriole proximal end (Dzhindzhev et al., 2014; Kratz et al., 2015; Moyer et al., 2015; Moyer and Holland, 2019). The newly assembled daughter centrioles are regulated at their distal ends, promoting centriole elongation and stability (Chen et al., 2002; Schmidt et al., 2009). The four resulting centrioles accumulate pericentriolar material (PCM) proteins, important for centriole maturation and microtubule (MT) nucleation (Dammermann et al., 2004; Gould and Borisy, 1977; Loncarek et al., 2008; Moritz et al., 1995). How centriole assembly proteins are regulated to ensure that only two centrioles are assembled in S phase remains elusive.

Centriole assembly is regulated to prevent the formation of multiple daughter centrioles (centriole overduplication), which can result in centrosome amplification (CA) (Denu et al., 2018; Li et al., 2014). CA leads to chromosome segregation defects, chromosome instability, aneuploidy, and tumorigenesis (Basto et al., 2008; Ganem et al., 2009; Levine et al., 2017; Marthiens et al., 2013; Salisbury et al., 2004). Centriole overduplication can be driven by aberrant transcript and protein levels of centriole assembly factors, such as overexpression of PLK4 (Coelho et al., 2015).

How centrosome assembly factors are regulated and become dysregulated at the transcript and protein level in diseases such as cancer, remains unclear. Emerging evidence suggests genes that encode centrosome proteins undergo transcriptional and post-transcriptional RNA metabolic control to regulate centriole duplication (Fan et al., 2015; Fischer et al., 2014; Ganapathi Sankaran et al., 2019; Ledoux et al., 2013; Lee et al., 2014; Phan et al., 2022; Stemm-Wolf et al., 2021; Zhang et al., 2019). Additionally, some mRNAs are locally translated at centrosomes to regulate centrosome protein levels. Only eight mRNAs are currently known to localize to centrosomes, implying that their encoded proteins may function in key centrosome-related processes (Safieddine et al., 2021).

One of these mRNAs is *CEP350*, which localizes to centrosomes during interphase and mitosis (Safieddine et al., 2021). CEP350 protein localizes to the distal ends of centrioles and regulates centriole assembly, elongation, and stability (Gonçalves et al., 2021; Karasu et al., 2022; Mojarad et al., 2017). CEP350 protein also promotes distal and subdistal appendage formation, further enabling MT organization (Huang et al., 2017; Karasu et al., 2022; Yan et al., 2006). Whether *CEP350* mRNA is locally translated at the centrosome to maintain CEP350 centrosome protein levels is unknown. Moreover, the mechanistic details localizing *CEP350* mRNA and protein to centrosomes is unclear, but like other centrosome-associated mRNAs, *CEP350* mRNA localization is co-translational, requiring active translation (Safieddine et al., 2021).

Centrosome mRNA localization is also coordinated by dynein, RNA binding proteins (RBPs), and centriolar satellites (Pachinger et al., 2025; Ryder et al., 2020; Safieddine et al., 2021; Sepulveda et al., 2018). Centriolar satellites are membraneless granules that cluster around centrosomes in a MT-dependent manner (Gheiratmand et al., 2019; Kubo et al., 1999; Ryu et al., 2024; Staples et al., 2012). Centriolar satellite proteins, like CEP131, interact with and promote the localization of centriole assembly proteins to regulate centriole duplication (Hall et al., 2023; Kodani et al., 2015; Lee and Stearns, 2013; Odabasi et al., 2019; Staples et al., 2012; Stemm-Wolf et al., 2021). Centriolar satellites also promote the localization of mRNAs that are important to maintain centrosome protein levels (Martinez et al., 2025; Pachinger et al., 2025). RBPs, including Unkempt (UNK), localize near CEP131 and bind centrosome-localized mRNAs, including *CEP350* (Arslanhan et al., 2020; Gheiratmand et al., 2019; Gupta et al., 2015; Martinez et al., 2025; Murn et al., 2016; Murn et al., 2015; Pachinger et al., 2025; Shah et al., 2024). Whether centriolar satellites directly promote the localization or metabolism of centrosome-localized mRNAs to support translation during centriole duplication remains unknown.

The regulation of centriole assembly proteins is primarily understood at the post-translational level (Balestra et al., 2021; Čajánek et al., 2015; Christian Wigley et al., 1999; Cunha-Ferreira et al., 2013; Fabunmi et al., 2000; Watanabe et al., 2016). To understand the transcript-level regulation of centriole assembly factors, we explored the localization of *CEP350* mRNA and protein to centrioles during S phase. CEP350 is required for PLK4-induced centriole overduplication but has limited effects on canonical centriole duplication. We find that *CEP350* mRNA is localized to centrosomes, in part, by the CEP131 centriolar satellite protein and the UNK RBP in a MT-dependent manner. Further, CEP131 and UNK promote *CEP350* mRNA levels that are important for maintaining CEP350 centrosome protein levels. CEP131 and UNK promote *CEP350* mRNA stability. The localization of *CEP350* mRNA is perturbed and sequestered by elevated levels of cytoplasmic CEP131 aggregates, but dispensable for CEP350 protein localization. CEP131, UNK, and CEP350 promote CA in triple negative MDA-MB-231 breast cancer cells, highlighting these proteins as potential therapeutic targets. In summary, CEP131 and UNK promote the localization of *CEP350* mRNA and protein to centrosomes, a process important for centriole overduplication.

## RESULTS

### CEP350 mRNA and protein localizes to centrosomes during S phase

To determine the role of CEP350 on centriole duplication, we depleted CEP350 using a small interfering RNA (siRNA), arrested RPE-1 cells in S phase, and assessed centriole assembly in endogenous and PLK4 overexpressed cells using the RPE-1-Tet-PLK4, Centrin2:GFP cell line, hereafter inducible-PLK4 cells (Hatch et al., 2010). The number of Centrin2:GFP-labeled centrioles was quantified to measure centriole duplication. A defect in canonical centriole duplication (no PLK4 induction) was defined by cells with fewer than the expected 4 centrioles. CEP350 depletion mildly increased centriole underduplication from 0% (siCONTROL) to 5% (siCEP350) (p = 0.002; Fig. S1, A) (Karasu et al., 2022). To test whether CEP350 impacts centriole overduplication, CEP350 was depleted in PLK4 induced cells. CEP350 depletion severely reduced centriole overduplication (Le Clech, 2008) (cells with more than four centrioles) from 80% (siCONTROL) to 20% (siCEP350) (p = 0.0003; Fig. 1, A). Therefore, CEP350 facilitates PLK4 induced centriole overduplication, but has limited effects on canonical centriole duplication.

**Figure 1.**
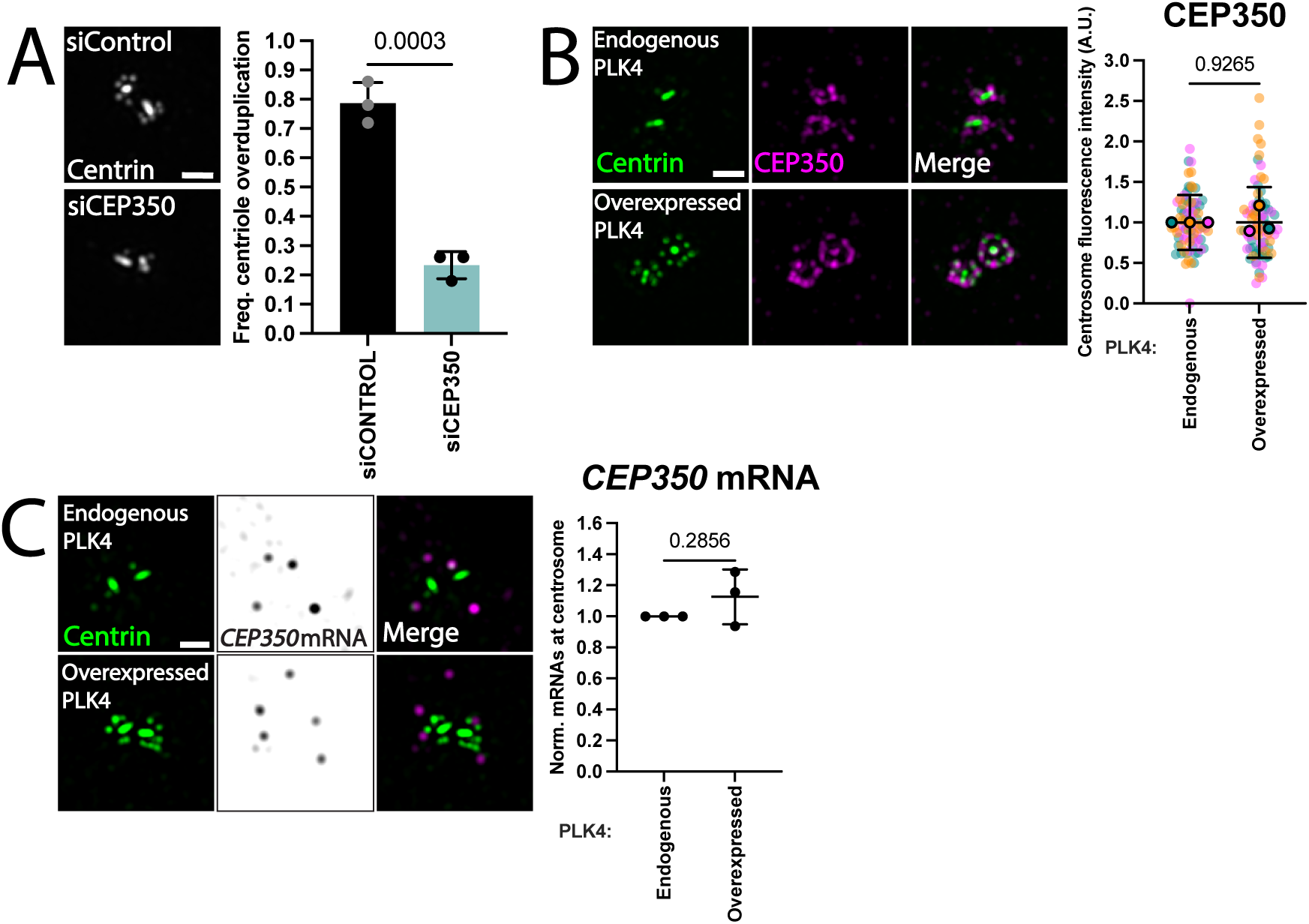
*CEP350* mRNA and protein localize to centrosomes in S phase. **(A)** CEP350 facilitates PLK4-induced centriole overduplication. Left panels: Structured illumination microscopy (SIM) images of CEP350-depleted RPE-1-Tet-PLK4, Centrin2:GFP cells showing centrioles with PLK4 induction in S phase. Centrioles (Centrin2:GFP), grayscale. Scale bar, 1.0 μm. Right panels: Frequency of RPE-1-Tet-PLK4, Centrin2:GFP cells with centriole overduplication in S phase. Graph values are expressed as the means of 3 biological replicates of 50 cells per replicate and SD. P values were determined using an unpaired two-tailed *t* test. **(B)** CEP350 localizes to centrioles in S phase. Left panels: SIM images of RPE-1-Tet-PLK4, Centrin2:GFP cells with endogenous and PLK4 induction in S phase. CEP350, magenta and centrioles (Centrin2:GFP), green. Scale bar, 1.0 μm. Right panels: Mean normalized centrosome fluorescence intensity of CEP350 protein. Graph values are expressed as the means of 3 biological replicates of 25-30 cells per replicate and SD. P values were determined using an unpaired 2 tailed *t* test. **(C)** *CEP350* mRNA localizes around centrosomes in S phase. Left panels: SIM images of RPE-1-Tet-PLK4, Centrin2:GFP cells showing centrioles and *CEP350* mRNA with endogenous and PLK4 induction in S phase. *CEP350* mRNA, grayscale and magenta, centrioles (Centrin2:GFP), green. Scale bar, 1.0 μm. Right panels: Quantification of the relative number of *CEP350* mRNAs at centrosomes. Graph values are expressed as the means of 3 biological replicates of 25-30 cells per replicate and SD. P values were determined using an unpaired 2 tailed *t* test.

To assess the localization of CEP350 protein during centriole assembly, we visualized endogenous CEP350 protein in inducible-PLK4 cells, with and without induction. Co-localization with Centrin2:GFP indicated that CEP350 protein localizes to both distal and proximal regions of mother and daughter centrioles in endogenous PLK4 cells (Fig. 1, B). Similarly, CEP350 protein localized to mother centrioles and overduplicated daughter centrioles in PLK4 induced cells (Fig. 1, B) (Karasu et al., 2022).

CEP350 protein levels at centrioles were quantified by immunofluorescence intensity using a 5 x 5 μm region of interest around both centrosomes, hereafter centrosomal levels, and compared between endogenous and PLK4 induced cells. CEP350 centrosomal protein levels were unchanged in PLK4 induced cells compared to endogenous PLK4 cells (Fig. 1, B). Thus, CEP350 protein localizes to mother and daughter centrioles in S phase, and the centrosomal levels of CEP350 are unchanged in cells with overduplicated centrioles. In summary, although CEP350 is required for centriole overduplication, total centrosomal CEP350 levels are similar between cells undergoing canonical centriole duplication and those with overduplicated centrioles.

### Centriolar satellites promote efficient localization, stability, and abundance of CEP350 mRNA

*CEP350* mRNA is one of few known centrosome-localized RNAs that localizes to centrosomes in mitosis and interphase (Safieddine et al., 2021). To determine if *CEP350* mRNA localizes to centrosomes during centriole duplication, we visualized the localization and quantified *CEP350* mRNAs using single molecule inexpensive fluorescence *in situ* hybridization (smiFISH) probes against endogenous *CEP350* mRNA, with and without PLK4 induction. *CEP350* mRNAs localize surrounding daughter centrioles (Fig. 1, C). The centrosomal *CEP350* mRNA levels modestly increased in PLK4 induced cells compared to endogenous PLK4 cells (p = 0.2856; Fig. 1, C and Fig. S1, B). In summary, *CEP350* mRNA localizes near daughter centrioles and *CEP350* mRNAs levels may be modestly increased in cells with overduplicating centrioles.

Centriolar satellites are granule structures that localize around centrosomes in a MT-dependent manner (Gheiratmand et al., 2019; Kubo et al., 1999). mRNAs and nascent peptides were previously shown to be in close proximity to centriolar satellites (Martinez et al., 2025; Pachinger et al., 2025). To determine whether *CEP350* mRNA localizes with centriolar satellites, we co-localized *CEP350* mRNA with the centriolar satellite protein CEP131 in inducible-PLK4 cells. *CEP350* mRNA localized closely to CEP131 puncta (Fig. 2, A). 60% of *CEP350* mRNA puncta localized within 0.5 μm of CEP131 protein puncta (associated), while 40% of *CEP350* mRNAs did not (alone) (Fig. 2, A). These data suggest that a majority of *CEP350* mRNA is closely associated with the centriolar satellite protein CEP131.

**Figure 2.**
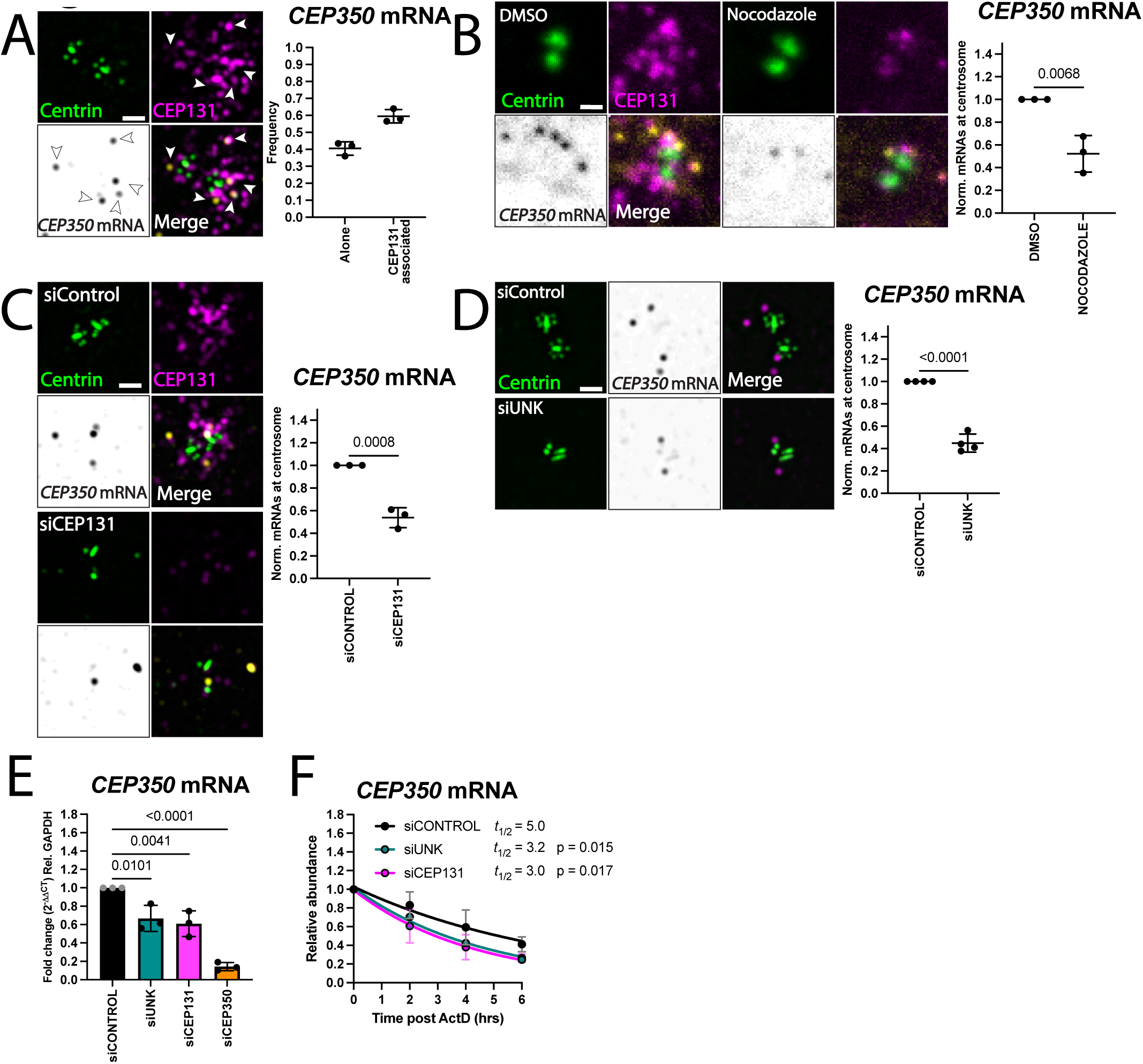
Centriolar satellites promote efficient localization and abundance of *CEP350* mRNA. **(A)** *CEP350* mRNA is closely associated with CEP131-positive centriolar satellites. Left panels: SIM images of RPE-1-Tet-PLK4, Centrin2:GFP cells showing centrioles, *CEP350* mRNA, and centriolar satellite protein CEP131 with PLK4 induction in S phase. *CEP350* mRNA, grayscale and yellow, CEP131, magenta, and centrioles (Centrin2:GFP), green. Scale bar, 1.0 μm. Right panels: Frequency of *CEP350* mRNAs alone and colocalized or within 0.5 μm of CEP131 (CEP131 associated). **(B)** MTs are required for efficient CEP131 and *CEP350* mRNA localization to centrosomes. Left panels: Confocal images of Nocodazole-treated RPE-1-Tet-PLK4, Centrin2:GFP cells showing centrioles, *CEP350* mRNA and centriolar satellite protein CEP131 with PLK4 induction in S phase. *CEP350* mRNA, grayscale and yellow, CEP131, magenta, and centrioles (Centrin2:GFP), green. Scale bar, 1.0 μm. Right panels: Quantification of the relative number of *CEP350* mRNAs at centrosomes. Graph values are expressed as the means of 3 biological replicates of 25-30 cells per replicate and SD. P values were determined using an unpaired two tailed *t* test. **(C)** CEP131 promotes efficient *CEP350* mRNA localization to centrosomes. Left panels: SIM images of CEP131-depleted RPE-1-Tet-PLK4, Centrin2:GFP cells showing centrioles, *CEP350* mRNA and centriolar satellite protein CEP131 with PLK4 induction in S phase. *CEP350* mRNA, grayscale and yellow, CEP131, magenta, and centrioles (Centrin2:GFP), green. Scale bar, 1.0 μm. Right panels: Quantification of the relative number of *CEP350* mRNAs at centrosomes. Graph values are expressed as the means of 3 biological replicates of 25-30 cells per replicate and SD. P values were determined using an unpaired 2 tailed *t* test. **(D)** UNK promotes efficient *CEP350* mRNA localization to centrosomes. Left panels: SIM images of UNK-depleted RPE-1-Tet-PLK4, Centrin2:GFP cells showing centrioles, *CEP350* mRNA and centriolar satellite protein CEP131 with PLK4 induction in S phase. *CEP350* mRNA, grayscale and magenta, and centrioles (Centrin2:GFP), green. Scale bar, 1.0 μm. Graph values are expressed as the means of 4 biological replicates of 25-30 cells per replicate and SD. P values were determined using an unpaired 2 tailed *t* test. **(E)** UNK and CEP131 promote *CEP350* mRNA steady-state levels. qRT-PCR quantification of *CEP350* mRNA total levels in UNK- and CEP131-depleted RPE-1-Tet-PLK4, Centrin2:GFP cells with PLK4 induction in S phase. *CEP350* mRNA levels were normalized to *GAPDH*. Graph values are expressed as the means of 3 biological replicates and SD. P values were determined using one-way ANOVA with Dunnett *post hoc* test. **(F)** CEP131 and UNK promote *CEP350* mRNA stability. qRT-PCR quantification of *CEP350* mRNA total levels in CEP131- and UNK-depleted RPE-1-Tet-PLK4, Centrin2:GFP cells with PLK4 induction in S phase, 2, 4 and 6 hrs after Act D treatment. *CEP350* mRNA levels were normalized to *RPS20* and expressed relative to time 0. Data were fit using a single-phase exponential decay model with the plateau constrained to 0. *CEP350* mRNA half-lives (t½) were calculated from the fitted decay constants. Decay rates were compared using an extra sum-of-squares F test. Data represent mean from 3 biological replicates and SD.

mRNAs use RBPs and molecular motors on MTs to localize to centrosomes (Ryder et al., 2020; Safieddine et al., 2021). To determine if *CEP350* mRNA localization to centrosomes is MT-dependent, we assessed the localization of *CEP350* mRNA after nocodazole depolymerization of MTs in inducible-PLK4 cells. Nocodazole treatment depolymerized microtubules (Fig. S1, C) and reduced CEP131 protein at centrosomes to 50% of control levels (Fig. 2, B and p = 0.002; Fig. S1, D; (Ryu et al., 2024)). Similarly, MT-depolymerization reduced centrosomal *CEP350* mRNA to 50% of control levels (p = 0.0068; Fig. 2, B and Fig. S1, D). These data suggest that *CEP350* mRNA and CEP131-positive centriolar satellites utilize MTs to localize to centrosomes. Half of *CEP350* mRNAs and CEP131-positive centriolar satellites remain at centrosomes upon MT depolymerization, possibly due to localization through microtubule-independent mechanisms, or due to the establishment of a stable association at centrosomes prior to microtubule depolymerization. Thus, centriolar satellites and *CEP350* mRNA localize around centrosomes, in part, by MTs.

Centriolar satellites promote the localization of mRNAs to centrosomes (Martinez et al., 2025; Pachinger et al., 2025). To test whether centrosome-associated *CEP350* mRNA is dependent on centriolar satellites, we assessed the localization of *CEP350* mRNA in CEP131 depleted, inducible-PLK4 cells. CEP131 depletion reduced CEP131-positive centriolar satellites at centrosomes and disrupted centriole overduplication (Fig. 2, C) (Kim et al., 2019; Martinez et al., 2025; Stemm-Wolf et al., 2021). Further, depletion of CEP131 reduced *CEP350* mRNAs at centrosomes to 50% of control levels (p = 0.0008; Fig. 2, C and p = 0.0001; S1, E). This suggests that CEP131 promotes the localization of *CEP350* mRNAs, but like conditions of MT depolymerization, half of *CEP350* mRNAs remain at centrosomes upon CEP131 loss. In summary, CEP131-positive centriolar satellites promote the efficient localization of *CEP350* mRNA around centrosomes.

### UNK RNA-binding protein promotes localization, stability, and abundance of CEP350 mRNA

The Unkempt (UNK) RBP localizes near centrioles and promotes the localization of CEP131-positive centriolar satellites (Martinez et al., 2025). UNK also binds centrosome-localized mRNAs, including *CEP350* mRNA (Murn et al., 2016; Murn et al., 2015; Shah et al., 2024) and promotes mRNA localization in neurons (Arora et al., 2022). To investigate whether UNK promotes centrosomal *CEP350* mRNA localization, we assessed *CEP350* mRNA localization in UNK depleted, inducible-PLK4 cells. UNK depletion decreased centriole overduplication (Fig. 2, D) (Martinez et al., 2025). UNK depletion also reduced centrosomal *CEP350* mRNAs to 50%, while centrosomal *CEP350* mRNA fluorescence intensity decreased 26% (p = <0.0001; Fig. 2, D and p = 0.0143; S1, F). This suggests that UNK promotes the efficient localization of *CEP350* mRNAs to centrosomes, but like conditions of MT depolymerization and CEP131 loss, some *CEP350* mRNAs remain at centrosomes upon UNK depletion.

Given that centriolar satellite loss reduces centrosomal *CEP350* mRNA, we asked whether this resulted from depletion of total *CEP350* mRNA levels. To establish whether UNK or CEP131 regulate steady state *CEP350* mRNA levels we measured *CEP350* transcripts by quantitative PCR (qPCR) in either UNK, CEP131, or CEP350 depleted, inducible-PLK4 cells. CEP350 depletion reduced *CEP350* mRNA to 10% of control levels (p = <0.0001; Fig. 2, E). UNK depletion reduced *CEP350* total mRNA levels to 62%, while CEP131 depletion reduced *CEP350* total mRNA levels to 60**%** (p = 0.0101 and 0.0041, respectively; Fig. 2, E). This suggests that UNK and CEP131 promote the steady-state levels of *CEP350* mRNA. Notably, this is in contrast to the centriolar satellite protein PCM1 which supports *PCNT* mRNA localization without affecting its mRNA levels (Pachinger et al., 2025). We propose that PCM1 and CEP131 have distinct roles in mRNA regulation. This is consistent with their opposing effects on centriole overduplication; PCM1 is dispensable for centriole overduplication, whereas CEP131 promotes it (Hall et al., 2023; Kim et al., 2019; Martinez et al., 2025; Stemm-Wolf et al., 2021). UNK and CEP131 may either regulate transcription of the *CEP350* gene (Shen et al., 2022), or stabilize *CEP350* mRNA. In summary, UNK and CEP131 promote *CEP350* mRNA localization to centrosomes and *CEP350* mRNA steady-state levels.

To determine whether UNK and CEP131 maintain *CEP350* mRNA steady-state levels by stabilizing *CEP350* mRNA, we inhibited transcription with Actinomycin D (Act D) and measured the *CEP350* mRNA rate of decay relative to the stable *RPS20* mRNA (Ratnadiwakara and Änkö, 2018; Spasic et al., 2012). UNK depletion reduced the half-life of *CEP350* mRNA from 5.0 hrs to 3.2 hrs (p = 0.015; Fig. 2, F). Similarly, CEP131 depletion reduced the half-life of *CEP350* mRNA to 3.0 hrs (p = 0.017; Fig. 2, F). These data suggest that both UNK and CEP131, two centriolar satellite proteins, stabilize *CEP350* mRNA.

### Overexpression of centriolar satellite protein CEP131 reduces centrosomal CEP350 mRNAs

Centriolar satellites can promote the localization of mRNAs and their encoded protein to centrosomes (Pachinger et al., 2025). We reasoned that loss of *CEP350* mRNAs observed in UNK and CEP131 depleted cells would decrease CEP350 centrosome protein levels. In support of this model, both UNK and CEP131 promote the localization of nascent polypeptides to centrosomes (Martinez et al., 2025). We quantified the centrosomal CEP350 protein in UNK and CEP131 depleted, incucible-PLK4 cells. CEP131 depletion modestly reduced CEP350 centrosomal protein levels to 80% of control levels (p = 0.0017; Fig. 3, A). UNK depletion reduced CEP350 centrosomal protein levels to 64% of control levels (p = 0.0129; Fig. 3, B). Thus, UNK promotes and has a stronger effect on centrosomal CEP350 protein localization than CEP131. The loss of CEP350 protein from the centrosome observed in UNK and CEP131 depleted cells could be attributed to decreased localized translation of *CEP350* mRNA. Because half of *CEP350* mRNAs remain at the centrosome in UNK and CEP131 depleted cells (Fig. 2, C and D), these remaining centrosomal *CEP350* mRNAs may support protein synthesis. Alternatively, the decrease in CEP350 protein observed in UNK- and CEP131-depleted cells may result from localization mechanisms that do not rely on local translation at centrosomes. In summary, these data support the model that UNK and CEP131 promote the efficient localization of CEP350 protein to centrosomes.

**Figure 3.**
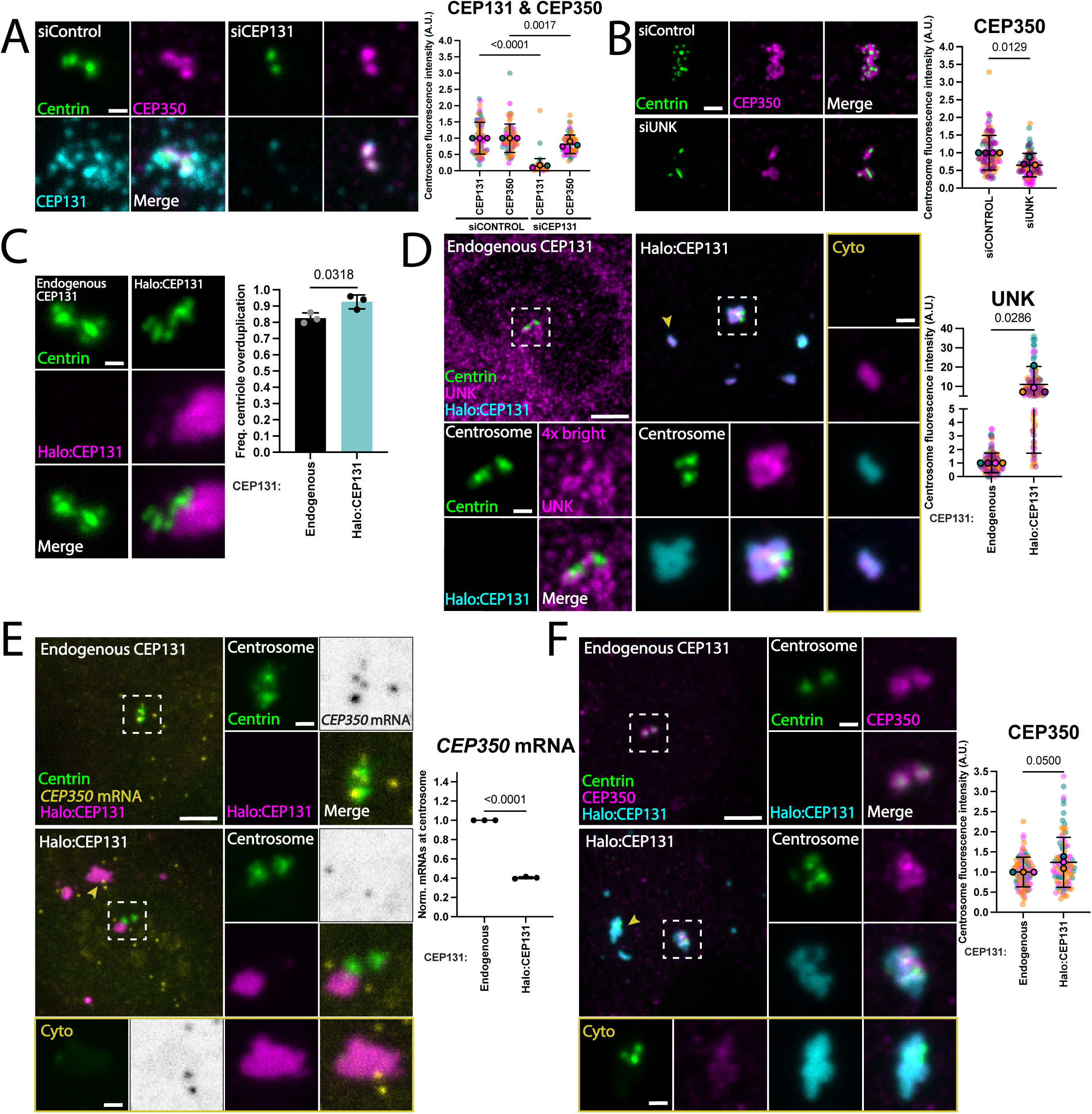
Overexpression of centriolar satellite protein CEP131 reduces centrosomal *CEP350* mRNA. **(A)** CEP131 promotes efficient localization of CEP350 protein to centrosomes. Left panels: Confocal images of CEP131-depleted RPE-1-Tet-PLK4, Centrin2:GFP cells showing centrioles, CEP350 protein and centriolar satellite protein CEP131 with PLK4 induction in S phase. CEP350 protein, magenta, CEP131, cyan, and centrioles (Centrin2:GFP), green. Scale bar, 1.0 μm. Mean normalized centrosome fluorescence intensity of CEP131 and CEP350 protein. Graph values are expressed as the means of 3 biological replicates and SD. P values were determined using one-way ANOVA with Šidák *post hoc* test. **(B)** UNK promotes efficient localization of CEP350 protein to centrosomes. Left panels: SIM images of UNK-depleted RPE-1-Tet-PLK4, Centrin2:GFP cells showing centrioles and CEP350 protein with PLK4 induction in S phase. CEP350 protein, magenta, and centrioles (Centrin2:GFP), green. Scale bar, 1.0 μm. Mean normalized centrosome fluorescence intensity of CEP350 protein. Graph values are expressed as the means of four biological replicates of 25-30 cells per replicate and SD. P values were determined using an unpaired 2 tailed *t* test. **(C)** CEP131 induction has modestly increased centriole overduplication. Left panels: Widefield images of RPE-1-Tet-PLK4, Centrin2:GFP cells showing centrioles and Halo:CEP131 with PLK4 and Halo:CEP131 induction in S phase. Halo:CEP131, magenta, and centrioles (Centrin2:GFP), green. Scale bar, 1.0 μm. Right panels: frequency of RPE-1-Tet-PLK4, Centrin2:GFP cells with centriole overduplication in S phase. Graph values are expressed as the means of 3 biological replicates of 50 cells per replicate and SD. P values were determined using an unpaired 2 tailed *t* test. **(D)** CEP131 induction promotes UNK localization to centrosomes. Left panels: Confocal images of RPE-1-Tet-PLK4, Centrin2:GFP cells showing centrioles, UNK, and Halo:CEP131 with PLK4 and Halo:CEP131 induction in S phase. UNK signal in endogenous CEP131 cells was displayed at 4X brightness relative to CEP131 overexpressing cells for visualization. UNK, magenta, Halo:CEP131, cyan, and centrioles (Centrin2:GFP), green. Scale bar, 5.0 μm. Insets scale bar, 1.0 μm. Right panels: Mean normalized centrosome fluorescence intensity of UNK. Graph values are expressed as the means of 4 biological replicates of 25-30 cells per replicate and SD. P values were determined using an unpaired 2 tailed *t* test. **(E)** CEP131 induction reduces *CEP350* mRNA from centrosomes. Left panels: Confocal images of RPE-1-Tet-PLK4, Centrin2:GFP cells showing centrioles, *CEP350* mRNA, and Halo:CEP131 with PLK4 and Halo:CEP131 induction in S phase. *CEP350* mRNA, grayscale and yellow, Halo:CEP131, magenta, and centrioles (Centrin2:GFP), green. Scale bar, 5.0 μm. Insets scale bar, 1.0 μm. Right panels: Quantification of the relative number of *CEP350* mRNAs at the centrosome. Graph values are expressed as the means of 3 biological replicates of 25-30 cells per replicate and SD. P values were determined using an unpaired 2 tailed *t* test. **(F)** CEP131 induction increases CEP350 centrosomal protein levels. Left panels: Confocal images of RPE-1-Tet-PLK4, Centrin2:GFP cells showing centrioles, CEP350 protein, and Halo:CEP131 with PLK4 and Halo:CEP131 induction in S phase. CEP350 protein, magenta, Halo:CEP131, cyan, and centrioles (Centrin2:GFP), green. Scale bar, 5.0 μm. Insets scale bar, 1.0 μm. Right panels: Mean normalized centrosome fluorescence intensity of CEP350 protein. Graph values are expressed as the means of 3 biological replicates of 25-30 cells per replicate and SD. P values were determined using an unpaired 2 tailed *t* test.

Because UNK and CEP131 promote centriole overduplication and promote the localization of *CEP350* mRNAs (Fig. 2, C and D), we asked whether overexpression of CEP131 impacts centriole overduplication and *CEP350* mRNA localization. A stable, doxycycline-inducible N-terminal HaloTag fusion to CEP131 (Tet-Halo:CEP131) was integrated into the inducible-PLK4 cells. When cells were induced for PLK4 and Halo:CEP131, centriole overduplication frequency increased from 80% (endogenous CEP131) to 92% (induced Halo:CEP131) (p = 0.318; Fig. 3, C). Halo:CEP131 localized to centrosomes, but also formed large cytoplasmic aggregates (Fig. S1, G). Centrosome and cytoplasmic aggregates co-localized with the PCM1 centriolar satellite protein, confirming they are characteristic of centriolar satellites. PCM1 co-localized with Halo:CEP131 around Centrin2:GFP labeled centrioles and the centrosomal levels of PCM1 protein were unchanged (p = 0.4835; Fig. S1, G). Moreover, both PCM1 and Centrin2:GFP co-localized with Halo:CEP131 in cytoplasmic aggregates (Fig. S1, G). This suggests that induced Halo:CEP131 aggregates that localize to centrosomes and the cytoplasm are centriolar satellites.

UNK and CEP131 are mutually dependent for their localization to centrosomes (Martinez et al., 2025). To determine whether induction of Halo:CEP131 is sufficient to drive UNK to centrosomes, inducible-PLK4 and Halo:CEP131 cells were stained for UNK protein. Strikingly, induction of Halo:CEP131 increased centrosomal UNK protein levels by approximately 10-fold relative to endogenous CEP131 cells (p = 0.0286; Fig. 3, D). Moreover, cytoplasmic aggregates formed by induction of Halo:CEP131 contained UNK protein (Fig. 3, D). These data are consistent with CEP131 promotion of UNK localization to centrosomes (Martinez et al., 2025). Whether Halo:CEP131 induction increases total UNK protein levels was not tested. In summary, induction of Halo:CEP131 is sufficient to increase UNK’s localization to centrosomes.

As centriolar satellite proteins UNK and CEP131 promote *CEP350* mRNA localization to centrosomes (Fig. 2 C and D), we hypothesized that induction of Halo:CEP131 and increased centrosomal UNK protein (Fig. 3, D) would increase *CEP350* mRNA localization to centrosomes. To test this, inducible-PLK4 and Halo:CEP131 cells were stained for *CEP350* mRNA. Unexpectedly, Halo:CEP131 induction reduced *CEP350* mRNAs at centrosomes to 40% compared to endogenous CEP131 cells (p = <0.0001; Fig. 3, E and S1, H). We hypothesize that cytoplasmic aggregates of Halo:CEP131 (Fig. 2, D and E, and S1, G) sequestered *CEP350* mRNAs from centrosomes. Consistent with this idea, *CEP350* mRNAs localized to the cytoplasmic Halo:CEP131 aggregates (Fig. 3, E). We conclude that induction of Halo:CEP131 reduces *CEP350* mRNAs at centrosomes by sequestering a limited pool of *CEP350* mRNAs to ectopic cytoplasmic aggregates.

Because induction of Halo:CEP131 reduced *CEP350* mRNAs from centrosomes, we asked whether this would impact CEP350 protein levels at centrosomes. Inducible-PLK4 and Halo:CEP131 cells were stained for endogenous CEP350 protein. Surprisingly, induction of Halo:CEP131 modestly increased centrosomal CEP350 protein levels by approximately 25% (p = 0.0500; Fig. 3, F). This indicates that induction of Halo:CEP131 reduces centrosomal *CEP350* mRNAs but does not impair CEP350 protein localization to centrosomes. This result differs from reduced centrosomal *CEP350* mRNA and corresponding reduced centrosomal CEP350 protein levels observed upon UNK and CEP131 depletion (Fig. 2, C and D; S1, E and F and Fig. 3, A and B). In summary, induction of Halo:CEP131 reduces *CEP350* mRNAs from centrosomes and modestly increases centrosomal CEP350 protein levels.

### CEP350 promotes centriolar satellite localization

Given that CEP350 promotes MT nucleation and organization (Fig. 4, A) (Yan et al., 2006), we asked whether CEP350 affects the MT-dependent localization of centriolar satellites (Ryu et al., 2024). CEP350 depleted, inducible-PLK4 cells were stained for CEP350 protein and the centriolar satellite protein CEP131. CEP350 depletion reduced centrosomal CEP350 protein levels by 70% (p = <0.0001; Fig. 4, B) and centrosomal CEP131 protein levels by 78% (p = 0.001; Fig. 4, B). The loss of CEP131 from the centrosome could be explained by the defective centrosomal MT nucleation and organization that is observed upon CEP350 loss (Fig. 4, A). Given that centriolar satellites also promote the localization of CEP350 to centrosomes (Fig. 3, A and B), these data suggest that CEP350 participates in a positive feedback loop that promotes the localization of centriolar satellites which in turn promote the localization of CEP350 to centrosomes.

**Figure 4.**
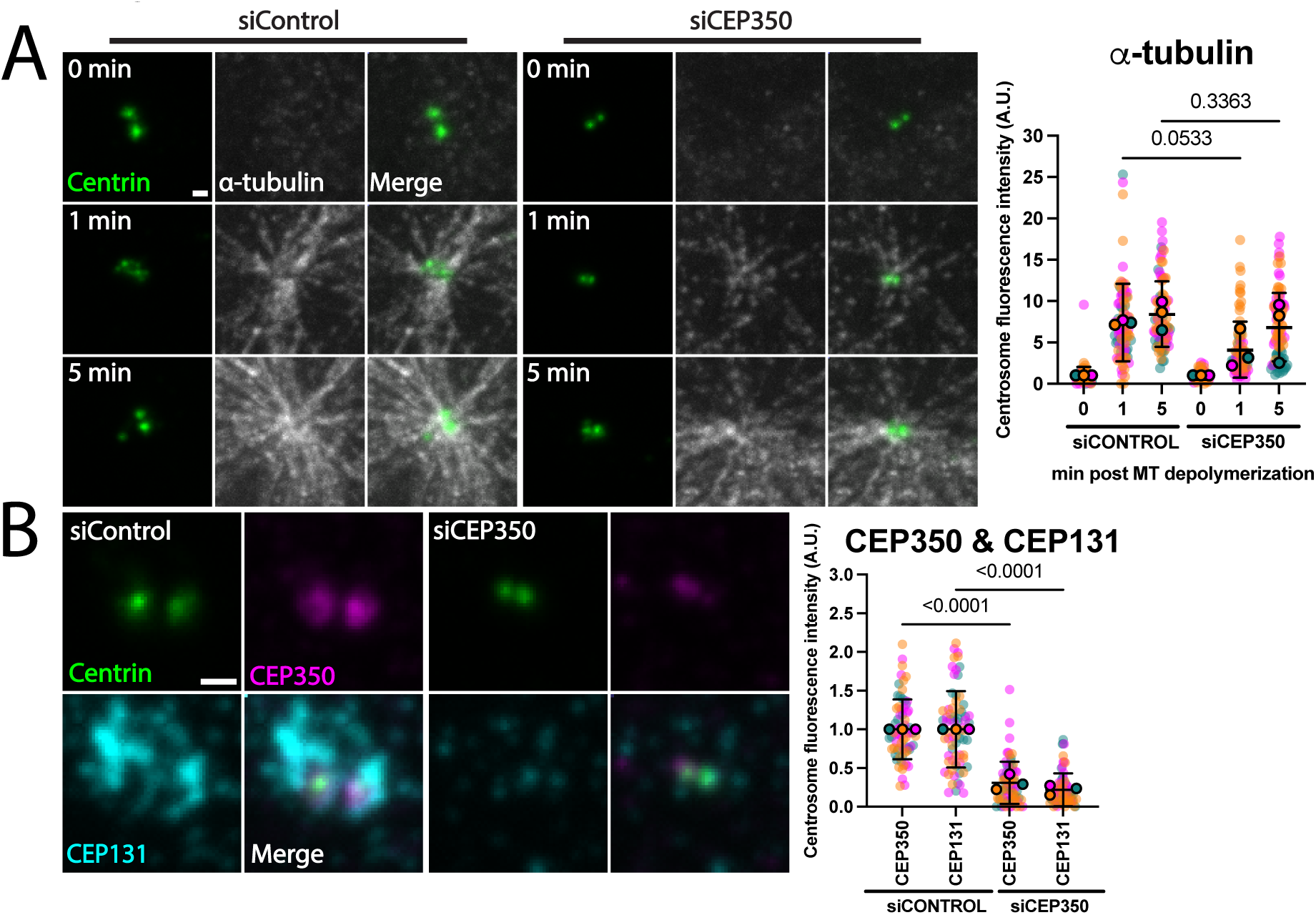
CEP350 promotes MT nucleation and centriolar satellite localization. **(A)** CEP350 promotes MT nucleation. Left panels: Confocal images of CEP350-depleted RPE-1-Tet-PLK4, Centrin2:GFP cells showing centrioles and α-tubulin 1 and 5 min, after depolymerization with PLK4 induction in S phase. α-tubulin, grayscale, and centrioles (Centrin2:GFP), green. Scale bar, 1.0 μm. Right panels: Mean normalized centrosome fluorescence intensity of α-tubulin. Graph values are expressed as the means of 3 biological replicates and SD. P values were determined using one-way ANOVA with Šidák *post hoc* test. **(B)** CEP350 promotes centriolar satellite localization to centrosomes. Left panels: Confocal images of CEP350-depleted RPE-1-Tet-PLK4, Centrin2:GFP cells showing centrioles, CEP350 protein, and centriolar satellite protein CEP131 with PLK4 induction in S phase. CEP350 protein, magenta, CEP131, cyan, and centrioles (Centrin2:GFP), green. Scale bar, 1.0 μm. Right panels: Mean normalized centrosome fluorescence intensity of CEP350 and CEP131 protein. Graph values are expressed as the means of 3 biological replicates and SD. P values were determined using one-way ANOVA with Šidák *post hoc* test.

### CEP350 facilitates centriole overduplication in triple negative breast cancer cells

Centrosome amplification (CA) is a common characteristic of cancer cells (Mittal et al., 2021) and PLK4-driven CA is sufficient to induce tumorigenesis in flies and mammals (Basto et al., 2008; Levine et al., 2017). Centriolar satellite proteins CEP131 and UNK facilitate PLK4-induced centriole overduplication but are dispensable for canonical centriole duplication (Hall et al., 2013; Martinez et al., 2025). Similarly, CEP350 supports centriole overduplication yet is not required for canonical centriole duplication (Fig. 1, A and S1, A). We asked if UNK, CEP131, or CEP350 depletion can reduce centriole overduplication in the triple negative breast cancer cell line MDA-MB-231 that commonly exhibits CA. To test this, MDA-MB-231 cells were depleted of UNK, CEP131, or CEP350, arrested in S phase, and assessed for centriole overduplication. 20% of siCONTROL treated MDA-MB-231 cells exhibited overduplicated centrioles (Fig. 5, A (D’Assoro et al., 2004; Ganapathi Sankaran et al., 2019; Kwon et al., 2008)). Depletion of UNK, CEP131, and CEP350 reduced centriole overduplication to 10%, 8%, and 7%, respectively (p = 0.0053, 0.0016, and 0.0012, respectively Fig. 5, A). This suggests that depletion of UNK, CEP131, or CEP350 suppresses centriole overduplication in triple negative breast cancer MDA-MB-231 cells.

**Figure 5.**
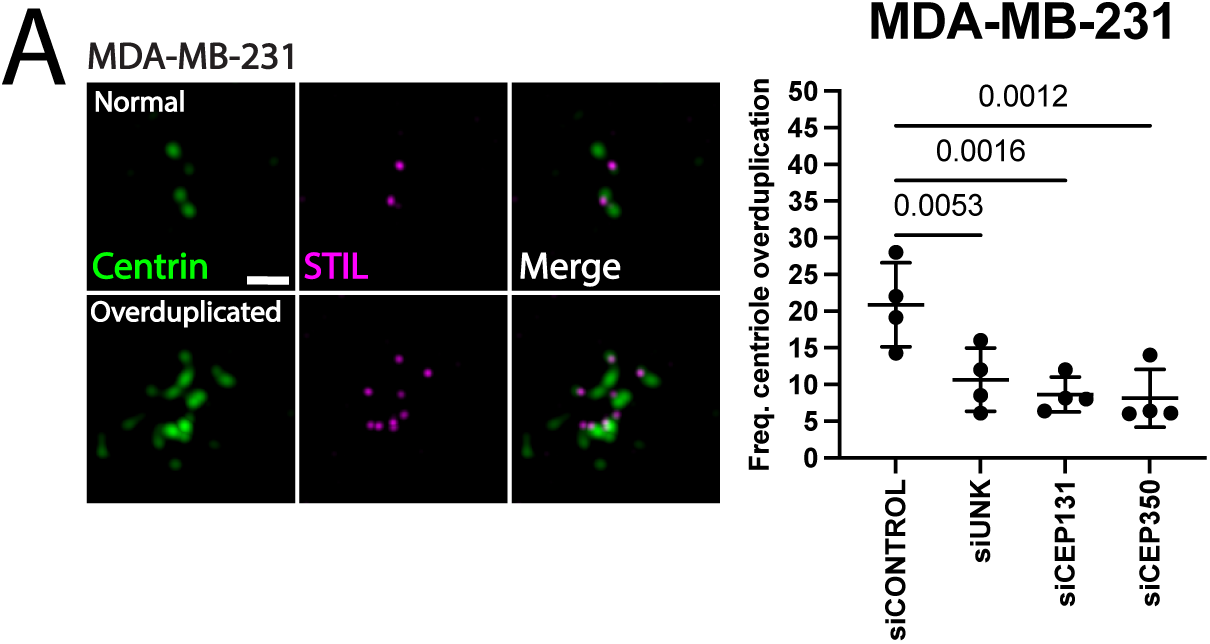
CEP350 facilitates centriole overduplication in triple negative breast cancer. **(A)** Left panels: SIM images of MDA-MB-231 cells showing centrioles and STIL protein in S phase. STIL, magenta, and centrioles (Centrin2:GFP), green. Scale bar, 1.0 μm. Right panels: Frequency of MDA-MB-231 cells with centriole overduplication in S phase. Graph values are expressed as the means of 4 biological replicates of 50 cells per replicate and SD. P values were determined using a one-way ANOVA with Fisher’s Least Significant Difference (LSD) *post hoc* test.

To assess the impact of UNK, CEP131, and CEP350 depletion on canonical centriole duplication, MCF10A immortalized mammary breast cells were depleted of either UNK, CEP131, or CEP350, S phase arrested and assessed for centriole underduplication. A loss in canonical centriole duplication was identified as cells with less than 4 centrioles. 4% of siCONTROL treated MCF10A cells had underduplicated centrioles (Fig. S1, I). Similarly, cells depleted of UNK and CEP131 had 2% centriole underduplication (p = 0.6115 and 0.5060, respectively; Fig. S1, I). CEP350 depletion increased centriole underduplication to 12% (p = 0.0320; Fig. S1, I). UNK and CEP131 have no detectable effects on canonical centriole duplication but significant effects on centriole overduplication, and may be favorable as cancer therapeutic targets to limit CA.

## DISCUSSION

Localized translation is a fundamental process that requires mRNA localization to subcellular structures and is utilized by differentiated and somatic cycling cells. In this study, we discovered that *CEP350* mRNA, one of few known centrosome-localized mRNAs, localizes closely with the centriolar satellite protein CEP131 during centriole duplication. The localization of *CEP350* mRNA to centrosomes is coordinated by the RBP UNK and the centriolar satellite scaffold protein CEP131 in a MT-dependent manner. Both UNK and CEP131 promote efficient localization of CEP350 protein to centrosomes and, in turn, CEP350 promotes CEP131 localization. CEP350 facilitates centriole overduplication. We reveal novel roles of centriolar satellites in regulating mRNA localization to centrosomes by positively regulating mRNA localization and stability.

We find that *CEP350* mRNA co-localizes with CEP131-positive centriolar satellites. Centriolar satellites are granular membrane-less structures that surround centrosomes (Aydin et al., 2020; Kodani et al., 2015; Kubo et al., 1999). Centriolar satellites and RBPs are thought to be phase condensates (Jiang et al., 2021; So et al., 2019). Phase condensates have RNA-regulatory roles including mRNA scaffolding and translation (Garcia-Jove Navarro et al., 2019; Jiang et al., 2021; Lin et al., 2015; Shapiro et al., 2025; So et al., 2019). Whether CEP131 or UNK transport, scaffold, and/or promote the localized translation of *CEP350* mRNA at centrosomes requires further investigation. This may also be mediated by the vast number of proteins that interact with centriolar satellites, including the many proteins involved in mRNA metabolic pathways (Gheiratmand et al., 2019; Quarantotti et al., 2019). Despite a loss of *CEP350* mRNA from centrosomes in UNK and CEP131 depleted cells, a subset *CEP350* mRNAs remain localized to centrosomes (Fig. 2, C and D; S1, E and F). It is possible only a specific subpopulation of centrosomal *CEP350* mRNAs relies on centriolar satellites for localization. Alternatively, once mRNAs are stably localized at centrosomes, they may be anchored to the pericentriolar material (PCM) (Fang et al., 2025).

Both UNK and CEP131 depletion result in similar levels of *CEP350* mRNA loss from centrosomes. However, the decrease in CEP350 protein at centrosomes was stronger upon UNK depletion than CEP131 depletion (Fig. 3, A and B). One possibility is that UNK, as a translation regulator, positively regulates *CEP350* local translation (Murn et al., 2016; Murn et al., 2015; Shah et al., 2024). In support of this model, we found that induction of CEP131 reduces *CEP350* mRNAs from centrosomes but has limited effects on centrosomal CEP350 protein levels (Fig. 3, E and F; S1, H). CEP131 induction drives increased centrosomal UNK levels (Fig. 3, D), which in turn, may promote local translation of centrosomal *CEP350* mRNAs. CEP131 induction also leads to the formation of cytoplasmic aggregates that sequester *CEP350* mRNAs away from centrosomes. These sequestered mRNAs may be ectopically translated into protein and moved to centrosomes (Fig. S1, G). Additionally, CEP131 and UNK have a role in stabilizing *CEP350* mRNA, which may contribute to normal CEP350 centrosomal protein levels upon CEP131 induction, even though centrosomal *CEP350* mRNA is reduced.

Beyond stabilizing and elongating centrioles, CEP350 promotes subdistal appendage formation, which anchor microtubules in interphase (Gonçalves et al., 2021; Huang et al., 2017; Karasu et al., 2022). Centriolar satellites and centrosomal proteins utilize MTs to localize to centrosomes (Fig. 2, B (Dammermann and Merdes, 2002; Ryu et al., 2024)). We found that CEP350 promotes centrosome MT nucleation (Fig. 4, A (Yan et al., 2006)) and is required for centriolar satellite localization. Thus, CEP350 loss impairs centriole overduplication both by its own absence and by preventing other molecules from localizing to centrosomes because of their reliance on MTs. Consistent with this, CEP350 loss has the strongest effect on centriole overduplication compared to CEP131 and UNK (Martinez et al., 2025).

Many cancer cell types are characterized by centriole overduplication and elevated *PLK4* expression (Ganapathi Sankaran et al., 2019; Mason et al., 2014). In addition to tumorigenesis, CA can result in epithelial to mesenchymal transition (EMT), and increased cell motility and invasion (Das, 2023; Godinho et al., 2014; Luo et al., 2019; Prakash et al., 2023). Current cancer drugs targeting PLK4 and cells with centrosome amplification in advanced triple negative breast cancer have reached phase II of clinical trials (ClinicalTrails.gov, 2026). However, whether this therapy can offer better results than traditional toxic chemotherapies is unknown (Riaz et al., 2025). One potential reason for limited efficacy is that PLK4 inhibition inhibits canonical centriole duplication (Bettencourt-Dias et al., 2005). An alternative approach would be to specifically target CA. We find that UNK, CEP131, and CEP350 are part of a pathway that are important for centriole overduplication. UNK and CEP131, and to a lesser extent, CEP350, are potential therapeutic targets for cancer cells with CA. Their depletion in breast cancer cells reduces CA. Importantly, depletion of these proteins exhibit minimal effects on canonical centriole duplication.

## METHODS

### Cell culture

MDA-MB-231 (University of Colorado Cancer Center Tissue Culture Core), RPE-1-Tet-PLK4, Centrin2:GFP (Hatch et al., 2010), and HEK293T cells were cultured in DMEM/F12 + L-glutamine (Cytiva or Life Technologies), 10% tetracycline-free FBS (Peak Serum), with 1% penicillin and streptomycin (Life Technologies) at 37°C and 5% CO_2_. MCF10A cells were cultured in Ham’s F/12 + L-glutamine (Corning), 5% horse serum (16050122; Thermo Fisher Scientific), Human EGF Recombinant Protein 20 ng/ml (PHG0311; Thermo Fisher Scientific), hydrocortisone 500 ng/ml (H0888; Sigma-Aldrich), cholera toxin 100 ng/ml (C8052; Sigma-Aldrich), insulin 10 µg/ml (I1882; Sigma-Aldrich), with 1% penicillin and streptomycin (Life Technologies) at 37°C and 5% CO_2_. A PCR test for mycoplasma contamination was performed every 6 mo.

### siRNA treatments

siRNAs used were: Mission siRNA Universal Negative Control #1 SIC001 (Sigma-Aldrich) at 50 nM, CEP131: Stealth HSS146116; Thermo Fisher Scientific 5′-CAGAGUGCCAGGAAUGCGGCAGCCU-3′ at 50 nM, UNK: Stealth HSS150335; Thermo Fisher Scientific 5′-CCUGACCUCAGUGCCCUCCUCUGU-3′ at 50nM, CEP350: pool of 4 ON-TARGET plus siRNAs L-015290-01-0005 5′-GUGAUUGGGUCUCGGGAAA-3′, 5′-GAUGAUAGGCAGUCGAGAA-3′, 5′-GUAGAGAACUGUAUCGAGA-3′, 5′-AGAUCUAAGUCGUCAGUAA-3′ at 50nM.

For 48 and 72 hr knockdown, 12,500 cells and 7,000 cells were plated on 12 mm circular cover glass, no. 1.5 (Electron Microscopy Sciences), coated with collagen (C9791; Sigma-Aldrich) in 24 well plates on day 0. On day 1, cells were treated with siRNAs using the Lipofectamine RNAiMAX Transfection Reagent (Thermo Fisher Scientific) according to the manufacturer’s instructions. Cells were provided fresh medium 6 hr later. On day 2, 1 hr prior to fixation, cells were arrested with 1.5 µg/ml aphidicolin (Cayman Chemical Company). Where indicated, PLK4 or CEP131 was overexpressed with 1.0 µg/ml doxycycline (Sigma-Aldrich) 8 hr after initiation of the aphidicolin arrest.

### qRT-PCR

Total RNA was harvested from cells using the Zymo *Quick*-DNA/RNA kit (D7005). DNase I was incubated on columns for 15 min at RT prior to eluting RNA in DEPC-treated water. 500ng of RNA was used for cDNA synthesis using SuperScript IV (Thermo Fisher Scientific) according to the manufacturer’s instructions. 200 ng of cDNA was utilized for qPCR using the Luna Universal qPCR Master Mix (NEB) according to the manufacturer’s instructions. cDNA was amplified on the QuantStudio 3 system (Thermo Fisher Scientific) at the annealing temperature of 60 °C. Primers used for *CEP350:* Fwd: 5’-GGCTCCAATACCAGGTTCTAAGC-3’, Rev: 5’-CATCTGACAGTTGGCGCACC-3’; *GAPDH*: Fwd: 5’-CCATGAGAAGTATGACAACAGCC-3’, Rev: 5’-GGGTGCTAAGCAGTTGGTG-3’. The relative fold changes were calculated using the ΔΔCt method with *GAPDH* as the reference gene (Crespo et al., 2020).

### Actinomycin D RNA stability assay

Cells were incubated with Actinomycin D (A9415; Sigma-Aldrich) at 10 µg/ml. Total RNA was harvested from cells following 2, 4, and 6 hrs using the Zymo *Quick*-DNA/RNA kit (D7005). RNA was harvested and processed for qRT-PCR. cDNA was amplified on the QuantStudio 3 system (Thermo Fisher Scientific) at the annealing temperature of 60 °C. The relative *CEP350* fold changes over time were calculated using the ΔΔCt method with *RPS20* as the reference gene (Ratnadiwakara and Änkö, 2018; Spasic et al., 2012). Primers used for *RPS20*: Fwd: 5’- AACAAGCCGCAACGTAAAATC-3’, Rev: 5’- ACGATCCCACGTCTTAGAACC -3’.

### Lentivirus transduction

Plasmid DNA was harvested using the ZymoPURE Plasmid Miniprep kit (D4208T). 2 μg of pCW57-HaloTag:CEP131, 1.5 μg of psPAX2 second-generation lentiviral packaging plasmid (12260; Addgene), and 0.5 μg of pMD2.G envelope-expressing plasmid in antibiotic-free Opti-MEM (31985070; Life Technologies), were transfected in HEK293T cells with using Lipofectamine 2000 Transfection Reagent (11668027; Thermo Fisher Scientific) according to the manufacturer’s instructions on day 0. On day 1, cells were replaced with fresh medium. On day 2 and 3, medium was removed from HEK293T cells, filtered, and added to RPE-1/Tet-PLK4, Centrin2:GFP with 10 µg/ml of polybrene. RPE-1-Tet-PLK4, Centrin2:GFP cells were selected with 10 µg/ml blasticidin for 1 week.

### Immunofluorescence

Cells on coverslips were washed one time with PBS and subsequently fixed with cold MeOH for 8 min at −20 °C. Coverslips were then washed three times in PBS, permeabilized with 0.5% Triton X-100 in PBS, and blocked in Knudsen buffer (1X PBS, 0.5% BSA, 0.5% NP-40, 1.0 mM MgCl_2_, and 1.0 mM NaN_3_) for 1 hr.

Cells stained for α tubulin (Fig. S1, C) were fixed with glutaraldehyde and paraformaldehyde (Canman et al., 2000). Briefly, cells were fixed with 4% paraformaldehyde and 0.5% glutaraldehyde in PIPES, HEPES, EGTA, and MGSO_4_ (PHEM) and permeabilized in 0.5% Triton X-100 in PHEM. Coverslips were carefully quenched three times with sodium borohydride in PHEM for 5 min each time, followed by 3, 5 min washes of 0.1% Triton X-100 PHEM and blocked with Knudsen buffer for 1 hr.

### MT regrowth assay

Cells were placed on ice and incubated at 4 °C for 20 min. Pre-warmed 37 °C media was placed on cells and incubated for 1 and 5 min. Cells were fixed with cold MeOH for 8 min at −20 °C. Coverslips were then washed three times in PBS, permeabilized with 0.5% Triton X-100 in PBS, and blocked in Knudsen buffer (1X PBS, 0.5% BSA, 0.5% NP-40, 1.0 mM MgCl_2_, and 1.0 mM NaN_3_) for 1 hr.

Antibody staining containing 1:1000 α-CEP350 (A301-170A; Bethyl) or 1:1,000 rabbit α-UNK (HPA023636; Sigma-Aldrich) antibodies were incubated overnight at 4°C only. All other antibodies were incubated for 1 hr at room temperature. Cells were washed four times for 5 min each in PBS prior to secondary antibody and DNA staining in Knudson buffer for 1 hr, except for cells stained for tubulin, in which fixation, staining, and washing were completed in PHEM buffer.

Primary antibodies: 1:1,000 rabbit α-STIL, and 1:1000 α-CEP192 (Moyer and Holland, 2019) (generous gifts from A. Holland, Department of Molecular Biology and Genetics, Johns Hopkins University School of Medicine, Baltimore, MD, USA), 1:5,000 guinea pig α-CEP131 (Kodani et al., 2015) (a generous gift from J. Reiter, Department of Biochemistry and Biophysics, University of California, San Francisco School of Medicine, San Francisco, CA, USA), 1:1,000 rabbit α-CEP131 (A301-415A; Bethyl), 1:2,500 rabbit α-CEP152 (A302-480A; Bethyl), and 1:500 mouse α-tubulin (CP06-100UG; Sigma-Aldrich).

Secondary staining: 1:1,000 Alexa anti-rabbit 488 and 594, 1:1,000 Alexa anti-mouse 647, and 1:1,000 Alexa anti-guinea pig 647, and 1:1,000 Alexa anti-goat 594 (Thermo Fisher Scientific). DNA was stained using Hoechst 33342 (62249; Thermo Fisher Scientific). Coverslips were mounted using Citifluor Af1 (17970-100; Electron Microscopy Services) and sealed with clear nail polish for confocal imaging and ProLong Gold Antifade (P10144; Thermo Fisher Scientific) for smiFISH and SIM.

### smiFISH

48 probes were generated using Stellaris probe design for the *CEP350* gene. DNA probes were ordered from IDT. The reverse complement of the X FLAP sequence, 5′-CCTCCTAAGTTTCGAGCTGGACTCAGTG-3′, was added to the 5′ end of each DNA probe. The probes were resuspended to a final concentration of 100 μM in TE buffer and then combined to make a final equimolar probe mix of 100 μM. To create smiFISH duplexes, a 10 μl mixture consisting of a final concentration of 20 μM equimolar probe mix, 1X NEB 3.1 buffer, and 25 μM Alexa Fluor 647-X FLAP (IDT) or iCy3-X FLAP (IDT), and DEPC-treated water was combined. The smiFISH duplexes were then assembled in a thermocycler at: 85°C for 3 min, 65°C for 3 min, 25°C for 5 min, and then kept on ice or frozen at −20°C for storage.

For HaloTag Janelia Fluor Dye labeling, cells were incubated with Janelia Fluor Dye 646 (Promega) in pre-warmed media for 45 min prior to fixation.

For RNA hybridization, cells were fixed onto 12 mm coverslips according to the LGC Biosearch Technologies RNA FISH adherent cells protocol. Briefly, cells were fixed with 3.7% formaldehyde in PBS for 10 min, followed by one PBS wash. 70% ethanol was used for permeabilization at 4°C for 1 hr or overnight. For each 12 mm coverslip, 0.5μL μl of smiFISH duplexes in 50 μl of Stellaris RNA FISH Hybridization buffer (SMF-HB1-10; LGC Biosearch Technologies) with 10% formamide was incubated on coverslips overnight at 37°C. The following day, coverslips were washed with Stellaris RNA FISH Wash Buffer A (SMF-WA1-60; LGC Biosearch Technologies) two times for 5 min. Coverslips were then placed in Stellaris RNA FISH Wash Buffer A for 30 min at 37°C, followed by one final wash in Stellaris RNA Wash Buffer B (SMF-WB1-20; LGC Biosearch Technologies).

For RNA hybridization with immunofluorescence, cells were fixed onto 12 mm coverslips according to the LGC Biosearch Technologies RNA FISH adherent cells + IF protocol. Briefly, cells were fixed with 3.7% formaldehyde in PBS for 10 min, followed by one PBS wash. 70% ethanol was used for permeabilization at 4°C for 1 hr or overnight. For each 12 mm coverslip, 0.5 μL of smiFISH duplexes in 50 μl of Stellaris RNA FISH Hybridization buffer with 10% formamide and the appropriately diluted antibody in DEPC-treated water was incubated on coverslips overnight at 37°C. The following day, coverslips were washed with Stellaris RNA FISH Wash Buffer A three times for 5 min. Coverslips were then placed in Stellaris RNA FISH Wash Buffer A with appropriately diluted secondary antibody for 30 min at 37°C. Coverslips were then washed three times for 5 min, followed by one final wash in Stellaris RNA Wash Buffer B.

### Fluorescence imaging

Confocal images were captured using a Yokogawa X1 spinning disk confocal on a Nikon Ti-E inverted microscope stand with a 100X Plan Apo NA 1.4 objective. Images were captured at room temperature using an Andor iXon EM-CCD camera with exposure settings between 0 and 500 msec and no binning of pixels using the SlideBook acquisition software.

Wide-field images were acquired with a Nikon Eclipse Ti-E microscope with a 100X Plan Apo NA 1.40 objective. Images were captured at room temperature using an Andor Xyla 4.2 scientific CMOS camera between 0 and 100 msec and no binning of pixels using the SlideBook acquisition software.

SIM images were captured using a Nikon SIM on a Nikon Ti2 (LU-N3-SIM; Nikon Instruments) microscope equipped with a 100X SR Apo TIRF, NA 1.49 objective. Images were captured using a Hamamatsu ORCA-Flash 4.0 Digital CMOS camera (C13440) with 0.1–0.2-μm Z step sizes. Exposure settings were between 0 and 200 msec, depending upon the experiment. All images were collected at 25°C using NIS Elements software (Nikon). Raw SIM images were reconstructed using the image slice reconstruction algorithm (NIS Elements).

### Centriole counts

Centrin2:GFP was used to count centrioles manually in RPE-1-Tet-PLK4, Centrin2:GFP cells, using the 100× PlanApo DIC, NA 1.4 objective with a 1.5X magnification optivar on a Nikon TiE inverted microscope stand. Fields of view were randomly chosen.

STIL, CEP192, and Centrin was used to count centrioles manually in MDA-MB-231 and MCF10A cells, using the 100X PlanApo DIC, NA 1.4 objective with a 1.5X magnification optivar on a Nikon TiE inverted microscope stand. Fields of view were randomly chosen.

### Image analysis

Image analysis was performed using FIJI (Schindelin et al., 2012). Image stacks were projected by maximum intensity.

Fluorescence intensities of centrosomes were measured within a 5 x 5 μm box encompassing both centrosomes. A measurement of cell intensity near both centrosomes was acquired for non-centrosome signal background subtraction. If subtraction of background signal generated a negative fluorescence signal, it was converted to 0. smiFISH puncta were manually counted within a 5 x 5 μm box encompassing both centrosomes.

### Statistical methods and data collection

Fluorescence intensity measurements between two groups were compared using an unpaired t test with two tails. Centriole counts between two groups were compared using a t-test with two tails. For multiple, or more than two comparisons, a one-way ANOVA with *post hoc* tests of Dunnett, Šidák, or Fisher’s least significant difference was conducted where indicated. Statistics were calculated using Prism (GraphPad). Investigators were not blinded when collecting data. Images were collected identically within experiments, and data analysis was automated to the extent possible to prevent bias.

## ACKNOWLEDGEMENTS

We are grateful to Alex Stemm-Wolf for feedback on the manuscript and present and past Pearson lab members who provided insights and comments on this project. Amber Baldwin and Neelanjan Mukherjee provided assistance with qRT-PCR experiments. Andrew Holland and Jeremy Reiter (UC San Francisco) generously shared antibodies. Meng-Fu Bryan Tsou (Sloan Kettering Cancer Center) supplied the RPE-1-Tet-PLK4, Centrin2:GFP cell line. This research was funded by NIH-NIGMS R35 GM140813 Diversity Supplement and W.M. Keck Foundation.

## Abbreviations

RBP: RNA binding protein
CA: Centrosome amplification
MT: Microtubule
smiFISH: single molecule inexpensive fluorescence in situ hybridization

---

**S. Figure 1.**
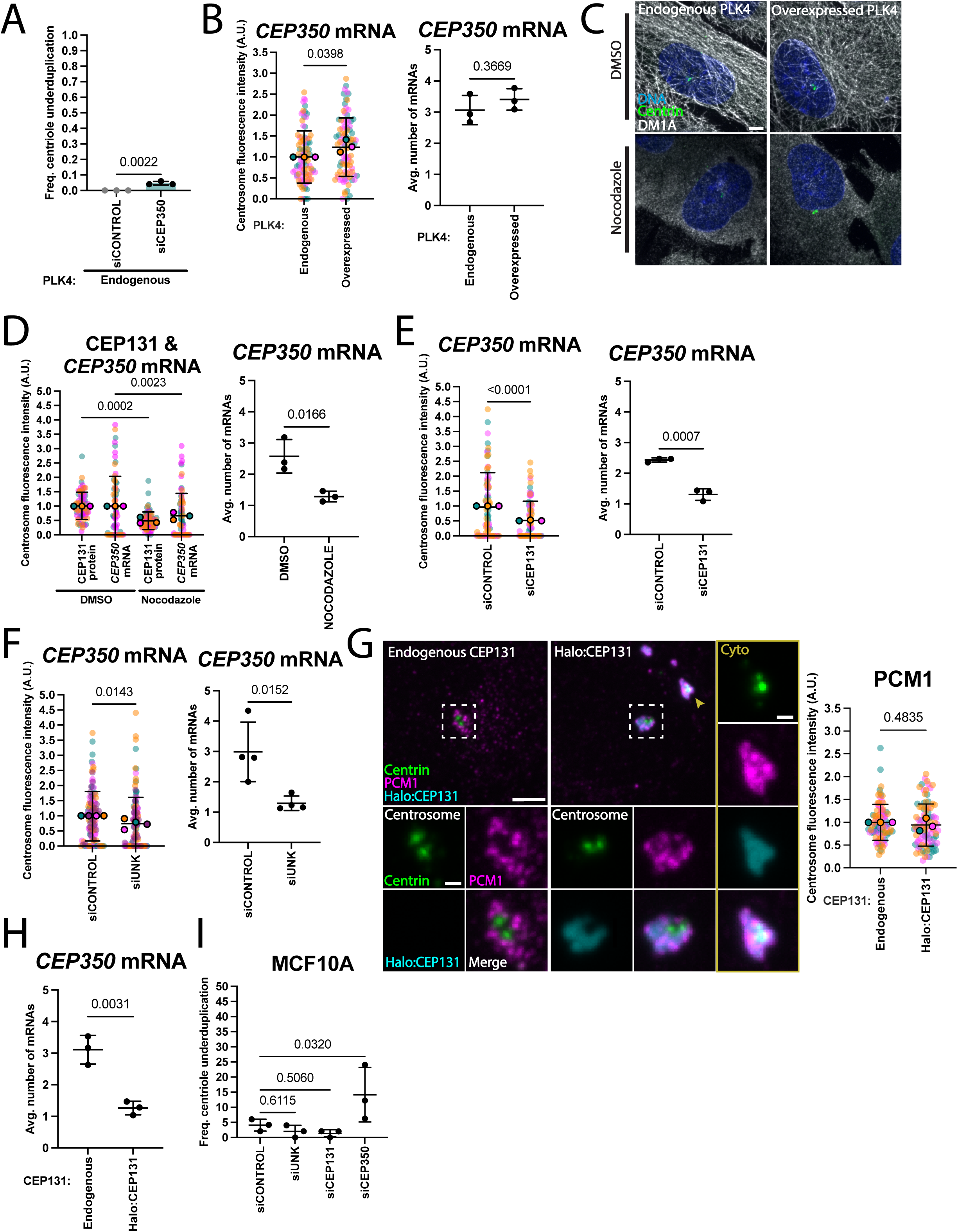
CEP350 facilitates PLK4 induced centriole overduplication. **(A)** CEP350 loss modestly increases centriole underduplication. Frequency of RPE-1-Tet-PLK4, Centrin2:GFP cells with centriole underduplication in S phase. Graph values are expressed as the means of 3 biological replicates of 50 cells per replicate and SD. P values were determined using an unpaired 2 tailed *t* test. **(B)** PLK4 induction has negligible effects on CEP350 centrosomal levels. Left panels: Mean normalized centrosome fluorescence intensity of *CEP350* mRNA. Graph values are expressed as the means of 3 biological replicates of 25-30 cells per replicate and SD. P values were determined using an unpaired two tailed *t* test. Right panels: Raw *CEP350* mRNA counts. Graph values are expressed as the means of 3 biological replicates of 25-30 cells per replicate and SD. P values were determined using an unpaired 2 tailed *t* test. **(C)** Nocodazole treatment depolymerizes MTs. Confocal images of nocodazole-treated RPE-1-Tet-PLK4, Centrin2:GFP cells showing centrioles, α-tubulin and DNA with endogenous and PLK4 induction in S phase. α-tubulin, grayscale, DNA, cyan, and centrioles (Centrin2:GFP), green. Scale bar, 5.0 μm. **(D)** MTs promote efficient localization of CEP131 protein and *CEP350* mRNA to centrosomes. Left panels: Mean normalized centrosome fluorescence intensity of CEP131 protein and *CEP350* mRNA. Graph values are expressed as the means of 3 biological replicates and SD. P values were determined using one-way ANOVA with Šidák *post hoc* test. Right panels: Raw *CEP350* mRNA counts. Graph values are expressed as the means of 3 biological replicates of 25-30 cells per replicate and SD. P values were determined using an unpaired 2 tailed *t* test. **(E)** CEP131 promotes efficient localization of *CEP350* mRNA to centrosomes. Left panels: Mean normalized centrosome fluorescence intensity of *CEP350* mRNA. Graph values are expressed as the means of 3 biological replicates of 25-30 cells per replicate and SD. P values were determined using an unpaired 2 tailed *t* test. Right panels: Raw *CEP350* mRNA counts. Graph values are expressed as the means of 3 biological replicates of 25-30 cells per replicate and SD. P values were determined using an unpaired 2 tailed *t* test. **(F)** UNK promotes efficient localization of *CEP350* mRNA to centrosomes. Left panels: Mean normalized centrosome fluorescence intensity of *CEP350* mRNA. Graph values are expressed as the means of 4 biological replicates of 25-30 cells per replicate and SD. P values were determined using an unpaired 2 tailed *t* test. Right panels: Raw *CEP350* mRNA counts. Graph values are expressed as the means of 4 biological replicates of 25-30 cells per replicate and SD. P values were determined using an unpaired 2 tailed *t* test. **(G)** CEP131 induction results in cytoplasmic aggregates. Left panels: Confocal images of CEP350-depleted RPE-1-Tet-PLK4, Centrin2:GFP cells showing centrioles, centriolar satellite protein PCM1, and Halo:CEP131 with PLK4 and halo:CEP131 induction in S phase. PCM1, magenta, Halo:CEP131, cyan, and centrioles (Centrin2:GFP), green. Scale bar, 5.0 μm. Insets scale bar, 1.0 μm. Right panels: Mean normalized centrosome fluorescence intensity of PCM1 protein. Graph values are expressed as the means of 3 biological replicates of 25-30 cells per replicate and SD. P values were determined using an unpaired 2 tailed *t* test. **(H)** CEP131 induction reduces *CEP350* mRNA from centrosomes. Raw *CEP350* mRNA counts. Graph values are expressed as the means of 3 biological replicates of 25-30 cells per replicate and SD. P values were determined using an unpaired 2 tailed *t* test. **(I)** UNK, CEP131, and CEP350 have minimal to modest effects on canonical centriole duplication. Graph values are expressed as the means of 3 biological replicates of 50 cells per replicate and SD. P values were determined using a one-way ANOVA with Fisher’s Least Significant Difference (LSD) *post hoc* test.

